# Dietary ethanol produces endpoint-specific dose responses in honeybees

**DOI:** 10.64898/2026.09.12.749844

**Authors:** Krzysztof Miler, Weronika Antoł, Monika Ostap-Chec, Daniel Bajorek

## Abstract

Ethanol is naturally present in nectar, but we still know little about how pollinators cope with long-term exposure to it at low concentrations. We exposed adult worker honeybees (*Apis mellifera carnica* Poll.) for eight days to diets containing 0, 0.1, 0.5, 1, 2, or 10% ethanol and measured survival, estimated intake, open-field locomotion, sting extension, whole-body trehalose, and hemolymph ethanol. Hemolymph ethanol was correlated with dietary concentration, attesting to internal exposure. Survival differed among diets, an effect which was driven by the difference between the 10% ethanol treatment and the control. Food intake was lowest at the 10% ethanol diet. There was also a modest increase in intake under the 0.1% ethanol treatment. In terms of locomotor features, bees moved less, moved more slowly, explored less space, covered shorter distances, and followed more tortuous paths after the 10% ethanol treatment. Neither sting extension nor trehalose were affected by dietary ethanol. The results showed no beneficial stimulation at lower concentrations. Instead, performance was endpoint-specific: at 0.1-2% ethanol in the diet, performance remained relatively stable across most parameters, whilst clear impairment was evident at 10%. These findings highlight resilience to repeated ethanol exposure in honeybees.

## Introduction

In response to stress, organisms often show what is known as hormesis or a biphasic dose-response (Mattson, 2008; Stebbing, 1982). A low amount of a stimulus improves some organismal parameters, but at higher intensities, the stimulus can have the opposite effect, inhibit performance, or become toxic. Hormesis has been reported amongst various taxa as a response to environmental factors such as temperature, water, or nutrition – a product of evolution to deal with stressors that organisms often have to face in nature (Agathokleous, 2018). At the cellular level, hormesis may be driven by signaling cascades that trigger adaptive stress responses including antioxidant systems, detoxification enzymes, and molecular chaperones (Mattson and Calabrese, 2010). These molecular processes can scale up to affect organismal performance and ecological interactions, positioning hormesis as a cross-level phenomenon bridging cellular physiology to behavior and life history trade-offs as well as ecological resilience (Agathokleous et al., 2018; Agathokleous et al., 2026). Importantly, however, a non-monotonic or curved response on its own does not indicate a clear-cut case of hormesis. At low doses of stressor, the organism has to not only perform at the same level as when it is not facing a challenge at all, but also show a more favorable outcome (Agathokleous, 2025). Additionally, hormetic effects are often modest, parameter-dependent, and constrained by organismal plasticity rather than producing large whole-organism benefits (Agathokleous et al., 2020; Calabrese et al., 2024). Low-dose stressors can also stimulate some traits while leaving others unchanged or even suppressing them (Erofeeva, 2023). Thus, any experiment looking at hormesis must cover a broad range of exposure levels and endpoints to resolve both beneficial and negative impacts across a wide dose gradient (Agathokleous, 2025). Identifying such relationships is essential for understanding the nature of hormesis and its physiological and/or ecological significance.

In insects, hormesis occurs in a variety of contexts but especially in agroecosystems, in which sublethal pesticide exposure can surprisingly increase reproduction, growth, longevity, and stress tolerance (Cutler et al., 2022). Upregulation of detoxification enzymes, heat shock proteins, and antioxidant activity that are commonly associated with these responses may be seen as manifestations of adaptive physiological plasticity, the ability to increase performance under mild environmental challenges (Rix and Cutler, 2022). Despite this, hormesis is underrepresented in pollinator ecotoxicology, where exposure effects are often split into binary safe/harmful categories (Cutler and Rix, 2015), even though sublethal stress might transiently enhance activity, foraging efficiency, or detoxification capacity in bees (Harwood et al., 2022; Strobl et al., 2020). Indeed, bees represent a relevant and interesting system for studying hormesis as their physiological responses can propagate across multiple biological levels. In social species, individual metabolic and neural responses dictate behavior, which in turn affects functional aspects like foraging performance and colony organization (Christen et al., 2019; Decourtye et al., 2003; Simone-Finstrom et al., 2022). Understanding when and how mild stress enhances performance is central to predicting pollinator resilience under environmental change.

Ethanol represents a particularly relevant natural stressor because it occurs in floral nectar and fermenting fruits, affecting a wide range of nectar- and fruit-feeding animals (Dudley, 2004; Dudley and Maro, 2021; Maro et al., 2026a). Such resources are consumed by many insects, which show clear behavioral and physiological adaptations to ethanol presence. For example, *Drosophila* species show well-documented tolerance to ethanol-rich substrates (Devineni and Heberlein, 2013; Starmer et al., 1977). In floral nectar, a recent survey discovered that ethanol occurs at approximately 0.008-0.056%, averaging 0.016% in general and 0.011% in honeybee-associated flowers (Maro et al., 2026a). Higher levels can be detected in fruits and some specialized systems (Dudley, 2004; Wiens et al., 2008). However, ethanol concentrations that occur in nature are well below those usually used to study the effects of ethanol on pollinators such as bees in the laboratory (Abramson et al., 2025).

Honeybees are known to accept ethanol-sucrose solutions across a broad range, and given the choice, they prefer them in some laboratory situations (Abramson et al., 2000; Abramson et al., 2004a; Mustard et al., 2019). However, intake or even preference is no evidence for benefits. In turn, concentrations of more than ∼5% clearly impact their motor skills, associative learning, and defensive behavior (Abramson et al., 2000; Black et al., 2021; Giannoni-Guzmán et al., 2014; Maze et al., 2006), and suppress adaptive stress responses (Hranitz et al., 2010). Despite this, they apparently do not form conditioned taste aversion to ethanol (Varnon et al., 2018). The effects of lower concentrations are less known. Acute exposure to 1% ethanol elevated hemolymph ethanol but did not detectably alter cognitive judgement bias or locomotor behavior in honeybees (Golański et al., 2026). Dietary ethanol at levels of 0.5% or 1% is generally less toxic to honeybees than higher concentrations (Ostap-Chec et al., 2024; Ostap-Chec et al., 2025a). Flight kinematics are altered but not impaired at 1% ethanol (Ahmed et al., 2022), and parasite-driven mortality decreases when honeybees have access to this concentration (Kuszewska, 2025; Ostap-Chec et al., 2025b). These findings raise the possibility that ethanol can produce beneficial responses in specific contexts. The overall dose-response relationship remains poorly resolved, however, because most studies have focused on relatively high concentrations within the toxic range and studies utilizing concentrations below 1% are almost absent (Abramson et al., 2025).

In this study, we exposed honeybees to dietary ethanol across a broad concentration gradient (0, 0.1, 0.5, 1, 2, and 10%). This series of concentrations linked lower, intermediate, and high-dose challenges. We measured survival and estimated food intake, open-field locomotor activity, aversive responsiveness in a sting extension test, and trehalose concentration as a metabolic indicator. To confirm differentiation among dietary treatments, we additionally measured ethanol levels in the hemolymph. We hypothesized that ethanol would stimulate physiological or behavioral performance at lower concentrations and become detrimental at higher concentrations.

## Materials and Methods

### Bee collection and maintenance

We used adult honeybees (*Apis mellifera carnica* Poll.) from three queen-right colonies headed by naturally inseminated queens. To obtain workers of broadly similar behavioral age, we adapted a previously described procedure (Miler et al., 2021). Each original hive was moved in the morning. A substitute hive was placed at its former location and supplied with food frames and uncapped and capped brood from the original colony, but no adult bees. Returning foragers entered the substitute hive, while many younger in-hive workers remained in the relocated colony. We collected workers that afternoon from randomly chosen frames in the relocated hives. Their exact age was unknown, but they were likely younger on average than active foragers, and so naïve to environmental ethanol.

We transported the bees to the laboratory and housed them in wooden cages, with 50 individuals per cage, in an incubator at 30 °C (KB400, Binder, Germany). Water and the assigned diet were available *ad libitum*. For each source colony, we prepared four sets of six treatment cages. The sets were used for four different assays: open-field behavior, sting extension, trehalose quantification, or hemolymph ethanol measurement. The full design comprised 72 cages (3 colonies × 4 assays × 6 diets), with 12 cages per dietary treatment and 3,600 bees at the start of exposure.

### Ethanol exposure treatments

Within each assay and source colony, we randomly assigned cages to 0, 0.1, 0.5, 1, 2, or 10% (v/v) ethanol treatments. We prepared the diets daily by adding absolute ethanol (99.8% v/v) to a 50% w/v aqueous Apikand solution (Łysoń, Poland). Each diet was mixed immediately before use and supplied in a gravity feeder. Exposure continued for eight days. We recorded mortality and feeder volume loss every day, then replenished the feeders with freshly prepared diet. At the end of the exposure period, bees were sampled for their respective assays.

### Open field assay

We measured locomotor performance in an open-field assay. Each bee was gently placed in the center of a circular Petri dish arena (14.5 cm diameter, 1 cm height) beneath a transparent lid. A top-mounted camera (Sony, Japan) recorded spontaneous activity for 10 min immediately after placement at 1920 × 1080 px and 50 fps. We used AnimalTA (Chiara and Kim, 2023) to reconstruct the trajectory of each bee. From these trajectories, we calculated the proportion of recording time spent moving, relative exploration (the proportion of arena area visited), mean speed while moving, total distance traveled, and meander (i.e., angular change per unit distance) used as an index of path tortuosity.

### Sting extension assay

We assessed aversive responsiveness by measuring sting extension during an ascending series of electric shocks. Each trial included seven stimuli, from 1 to 7 V in 1 V increments, delivered at 10 s intervals on an automated platform (Biospekt, Poland). A front-facing camera (Sony, Japan) recorded the 1.5-2 min sequence at 1920 × 1080 px and 50 fps. Using FFmpeg, we extracted frames at 5 fps (i.e., one frame every 200 ms) and imported them as image stacks into Fiji (Schindelin et al., 2012). We calibrated spatial scale separately for each video from an in-frame reference. For every stimulus, we measured the frame immediately before current onset and the frame with the greatest visible extension during current application. If no extension was visible, the response was recorded as 0 mm. Positive extension was measured as the straight-line distance from the sting emergence point to the visible tip. This gave 14 measurements per bee: seven before and seven during stimulation.

### Trehalose and hemolymph ethanol assays

Bees assigned to trehalose analysis were frozen immediately at -20 °C and stored individually until processing. We measured whole-body trehalose in 2 mL homogenates with a K-TREH assay kit (Neogen, USA), following the manufacturer’s protocol. For ethanol measurements, we cut one antenna and applied gentle pressure to the abdomen to collect hemolymph (Borsuk et al., 2017). The hemolymph was drawn into a 10 µL microcapillary, transferred to a cryotube, and stored at -20 °C. Samples containing at least 3 µL were brought to 10 µL with distilled water and analyzed with a K-ETOH assay kit (Neogen, USA), again following the manufacturer’s protocol. Each plate included standards spanning 0.00625-0.4 g/L for trehalose or 0.0125-0.1 g/L for ethanol. We measured the absorbance increase from the enzymatic reaction at 340 nm with a Multiskan FC microplate reader (Thermo Scientific, USA). Concentrations were calculated from plate-specific calibration curves and corrected for dilution.

### Statistics

We ran all analyses in R (R Core Team, 2025) and prepared figures with ‘ggplot2’ (Wickham, 2016). Diet was assigned at cage level, so the cage was used as the experimental unit to limit pseudo-replication issues. Within each assay, every source colony contributed one independently treated cage per diet, giving three cages per treatment. Mortality and feeder volume loss were recorded in all four assay-specific cage sets. These two analyses included 12 experimental runs (source colony × assay set), each with one cage from every dietary treatment. Estimated marginal means and 95% confidence intervals were obtained with ‘emmeans’ (Lenth, 2025) and back-transformed when needed. We used Dunnett adjustment for treatment-control comparisons and the Holm method to control multiplicity across the five locomotor endpoints. Heteroscedasticity-consistent (HC3) covariance estimates were calculated with ‘sandwich’ (Zeileis et al., 2026). Small-sample cluster-robust (CR2) tests were implemented with ‘clubSandwich’ (Pustejovsky, 2026). We checked model assumptions from residual plots and with ‘performance’ where appropriate (Lüdecke et al., 2021).

### Survival

For each cage, we calculated restricted mean survival through day 8 (RMST8) as the area under the daily survival-proportion curve and expressed it in days. Mortality was recorded once daily, so we assigned deaths within each 24-h interval to its midpoint. We analyzed RMST8 with a linear model containing dietary treatment and experimental run as fixed effects. We used HC3 covariance estimates. We first tested the overall treatment effect with a robust Wald F-test, then compared each treatment with the control using Dunnett adjustment.

### Food intake

For each cage, we calculated mean daily estimated intake over the eight-day exposure as total feeder volume loss divided by total estimated bee-days. We approximated bee-days within each 24-h interval from the mean numbers of bees alive at its beginning and end. This was equivalent to assigning deaths to the interval midpoint. We analyzed mean daily estimated intake with a linear model containing dietary treatment and experimental run as fixed effects. Here too we used HC3 covariance estimates. We first tested the overall treatment effect with a robust Wald F-test, then compared each treatment with the control using Dunnett adjustment.

### Locomotor behavior

Locomotor analyses used trajectories exported from AnimalTA. We excluded one video because tracking had failed. Almost all bees moved, so we only summarized movement initiation descriptively. Continuous traits were analyzed for bees that moved during the recording. We logit-transformed the proportion of time moving and relative exploration, and log-transformed mean speed while moving, distance traveled, and meander. Individual values were then averaged within cages to give one value per locomotor trait for each cage. We fitted linear models with dietary treatment and source colony as fixed effects. Overall treatment effects were tested with F-tests and adjusted across the five outcomes using the Holm method. For endpoints that retained an overall treatment effect after adjustment, we used Dunnett-adjusted contrasts to compare each ethanol treatment with the ethanol-free control.

### Sting extension

We excluded two individuals whose mounting did not allow reliable measurement. Sting extension during the ascending series of seven electric stimuli was used as the measure of aversive responsiveness. At each voltage step, we averaged measurements within treatment cages and retained zeros when no extension was visible. We analyzed these cage-level responses with a linear mixed model. Dietary treatment, voltage step, their interaction, and source colony were fixed effects. Voltage was entered as a continuous covariate centered at 4 V. Treatment cage had uncorrelated random intercepts and voltage slopes, which accounted for repeated measurements across the series and differences among cages in their response trajectories. We tested fixed effects with Satterthwaite-adjusted F-tests implemented in ‘lmerTest’ (Kuznetsova et al., 2017) and ‘lme4’ (Bates et al., 2015). Treatment-control contrasts used Dunnett adjustment. Pre-stimulus measurements were analyzed separately and analogically, as a supplementary control.

### Trehalose levels

We excluded one sample that had been identified as a technical outlier before analysis. Trehalose concentrations included a small number of exact zeros; the remaining values formed a right-skewed continuous distribution. We fitted a Tweedie generalized linear model with a log link. Dietary treatment, source colony, and assay plate were fixed effects. To account for dependence among bees from the same cage, we used CR2 covariance estimates clustered by cage. The overall treatment effect was tested with an approximate Hotelling T^2^ small-sample Wald test.

### Hemolymph ethanol concentration

Many hemolymph ethanol measurements were exactly zero, so we used a two-part analysis (Smith et al., 2014). In both components, dietary concentration was represented as log_10_[1 + concentration (%)/0.1]. This transformation retained the ethanol-free control while narrowing the spacing among higher concentrations. A binomial generalized linear model with a logit link estimated the probability of an exact zero. Positive concentrations were modeled separately with a Gamma generalized linear model and log link. Both components included source colony as a fixed effect. We accounted for dependence among bees from the same cage with CR2 covariance estimates clustered by cage. Dose effects were tested with approximate Hotelling T^2^ small-sample Wald tests.

## Results

### Survival

Of the 3,600 bees, 405 died during the exposure period. After accounting for experimental run, cage-level restricted mean survival time (RMST8) differed among dietary treatments overall (F_5,55_ = 3.16, p = 0.014; Fig. 1A). Estimates ranged from 7.568 days at 10% ethanol to 7.893 days at 1%. The largest difference from the control (7.856 days) occurred at 10% ethanol (difference = -0.288 days, 95% CI [-0.583, 0.008], Dunnett-adjusted p = 0.060). None of the other treatments differed from the control (all p ≥ 0.686; Supplementary Material 1: Table S1.1).

**Figure 1.**
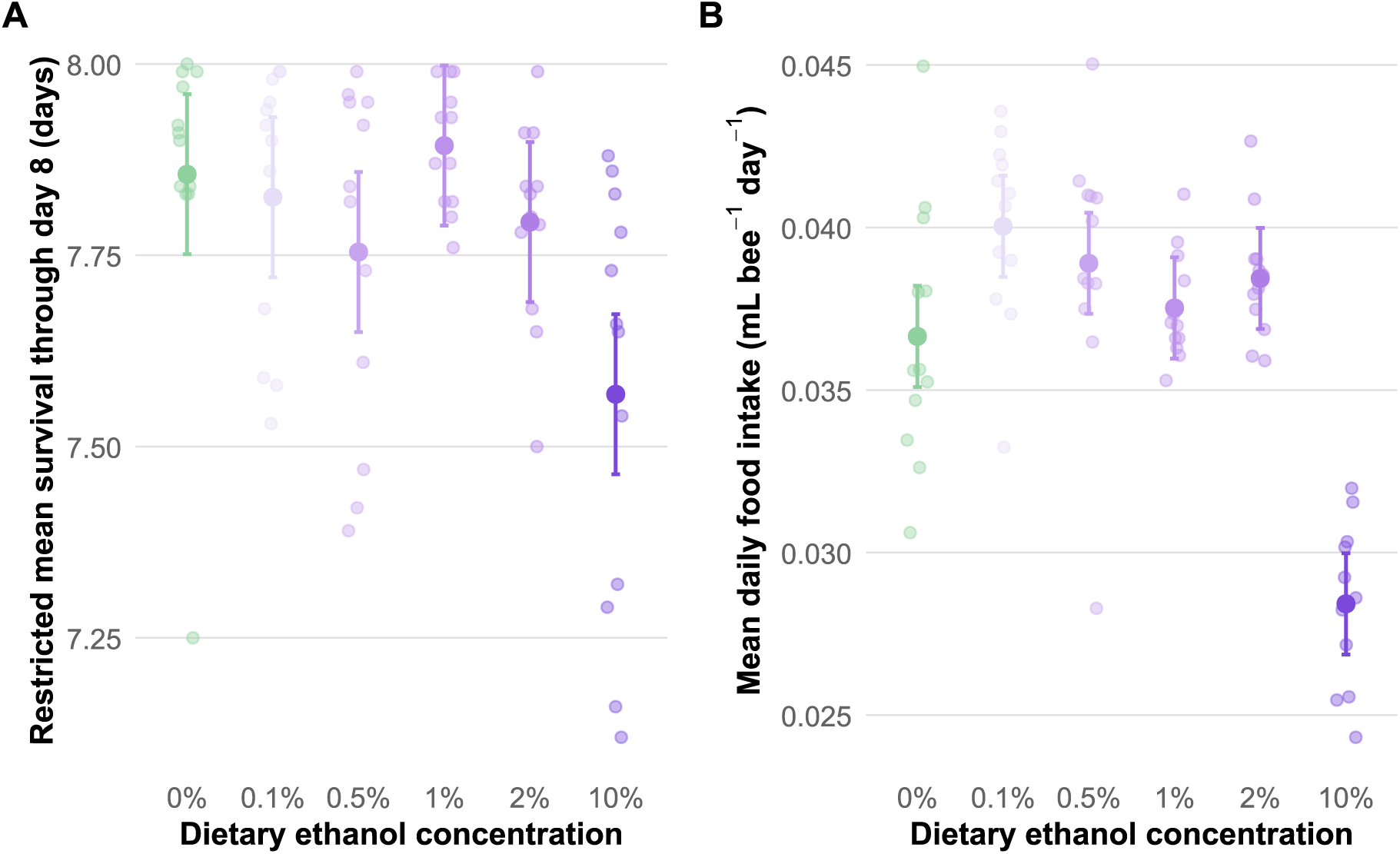
Effects of dietary ethanol on survival and estimated food intake. (A) Restricted mean survival through day 8. (B) Mean daily estimated intake per bee across the eight-day exposure. Semi-transparent points represent individual treatment cages. Larger opaque points and error bars show run-adjusted means and 95% confidence intervals from the cage-level models.

### Food intake

Mean daily estimated food intake differed among diets after accounting for experimental run (F_5,55_ = 16.88, p < 0.001; Fig. 1B). Intake was 0.0367 mL bee^−1^ day^−1^ in controls. The highest estimate, 0.0400 mL bee^−1^ day^−1^, occurred at 0.1% ethanol. The increase at 0.1% was 0.0034 mL bee^−1^ day^−1^ (approximately 9%), and it was borderline significant (95% CI: −0.0001 to 0.0068; Dunnett-adjusted p = 0.055). Intake was lowest at 10% ethanol (0.0284 mL bee^−1^ day^−1^). The reduction relative to controls was 0.0082 mL bee^−1^ day^−1^ (95% CI: −0.0121 to −0.0044; p < 0.001). Estimates at 0.5%, 1%, and 2% did not differ from the control (all p ≥ 0.357; Supplementary Material 1: Table S1.2).

### Locomotor behavior

Movement initiation was uniformly high: 324 of 332 bees moved during the recording, and the proportion of moving individuals ranged from 96.4% to 100% among treatments. Among these bees, dietary treatment affected the proportion of time moving (F_5,10_ = 7.84, p = 0.003), relative exploration (F_5,10_ = 8.36, p = 0.002), average speed while moving (F_5,10_ = 5.04, p = 0.015), distance traveled (F_5,10_ = 7.46, p = 0.004), and meander (F_5,10_ = 11.04, p < 0.001; Fig. 2), after accounting for source colony. All five effects remained significant after Holm adjustment (adjusted p ≤ 0.015; Supplementary Material 1: Table S1.3). The clearest changes occurred at 10% ethanol. Compared with controls, bees in this group spent less time moving (estimated proportion: 0.594 vs 0.788; Dunnett-adjusted p = 0.006), explored less (0.452 vs 0.741; p = 0.001), moved more slowly (2.279 vs 2.686 cm/s; p = 0.030), and traveled shorter distances (599 vs 1062 cm; p = 0.008). Meander increased from 333 deg/cm in controls to 421 deg/cm at 10% (p = 0.001). It was also elevated at 2% ethanol (391 deg/cm; p = 0.018). No other significant contrasts were detected. Although the 0.5% treatment showed higher estimates for several measures of locomotor performance, these differences were not statistically supported (Supplementary Material 1: Table S1.3).

**Figure 2.**
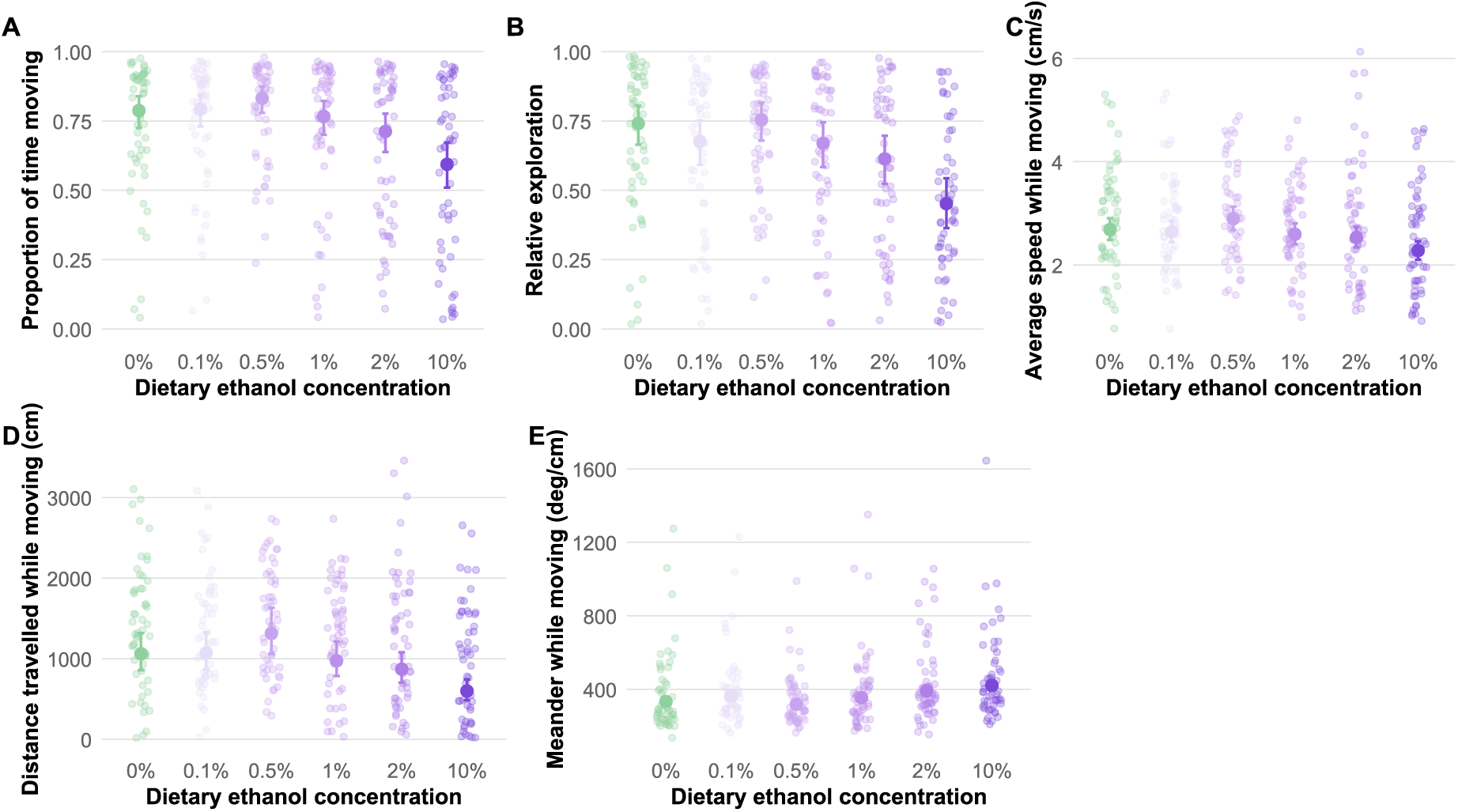
Locomotor activity across dietary ethanol treatments. Panels show (A) proportion of time moving, (B) relative exploration, (C) average speed while moving, (D) distance traveled while moving, and (E) meander while moving. Semi-transparent points represent individual bees that moved during the assay. Larger opaque points and error bars show source-colony-adjusted estimates and 95% confidence intervals from the cage-level models, back-transformed to the response scale.

### Sting extension

Sting extension increased across the ascending voltage series (F_1,12_ = 142.54, p < 0.001; Fig. 3). We found no evidence of an overall dietary treatment effect (F_5,10_ = 1.62, p = 0.241) or a treatment × voltage interaction (F_5,12_ = 0.54, p = 0.740). Voltage-response slopes ranged from 0.080 to 0.123 mm V^−1^, and none differed from the control after Dunnett adjustment (all p ≥ 0.841; Supplementary Material 1: Table S1.4). At the midpoint of the series, model-estimated sting extension ranged from 0.656 to 0.864 mm among ethanol treatments and was 0.741 mm in controls. Again, no treatment differed from the control (all Dunnett-adjusted p ≥ 0.529; Supplementary Material 1: Table S1.4). Chronic dietary ethanol did not detectably change either the overall magnitude of sting extension or its increase across the series (Supplementary Material 1: Table S1.5). Pre-stimulus sting extension also did not differ among diets (F_5,10_ = 1.05, p = 0.439; Supplementary Material 1: Table S1.6).

**Figure 3.**
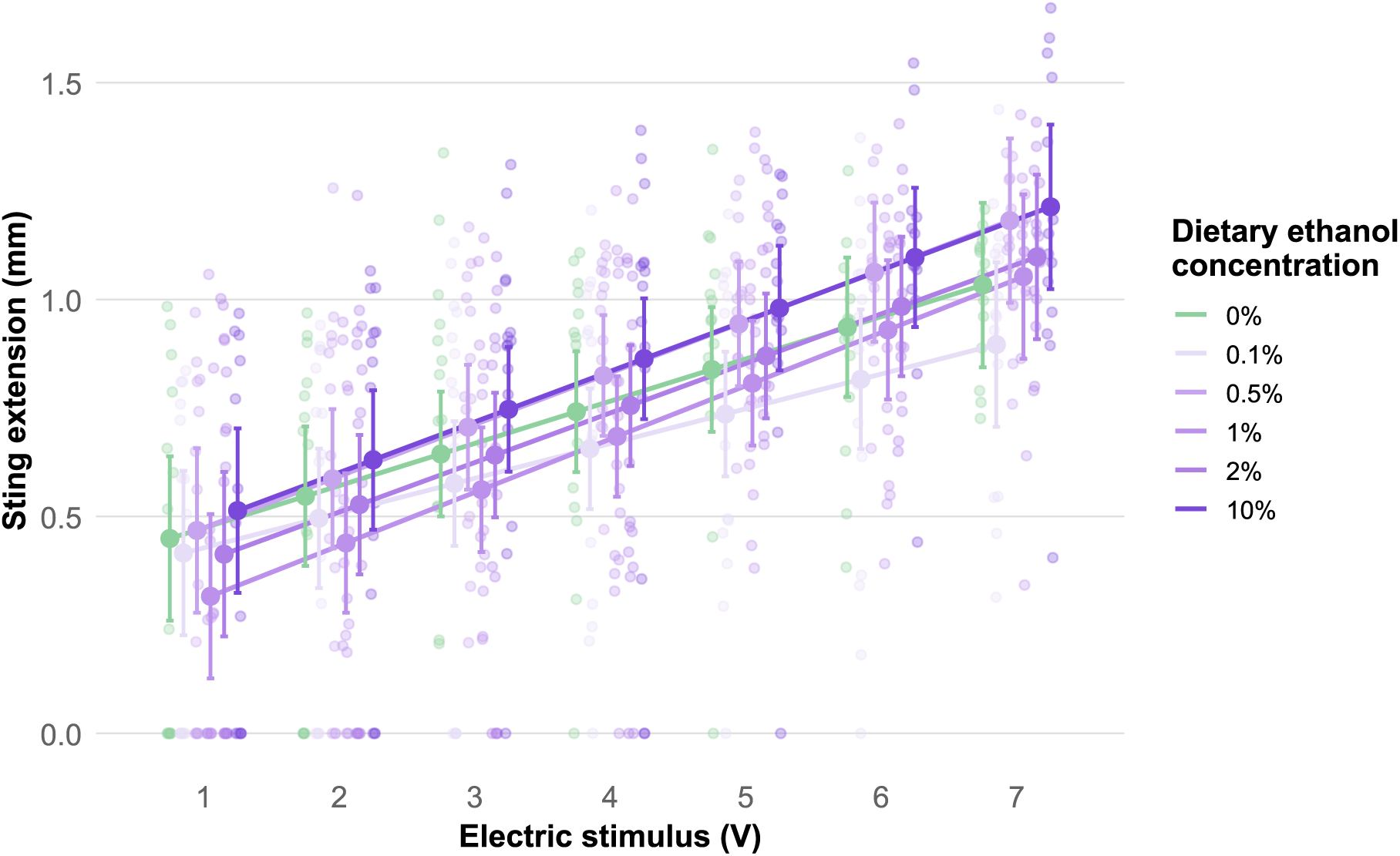
Sting extension during electric stimulation across dietary ethanol treatments. Semi-transparent points represent individual observations, including zero responses. Larger opaque points and error bars show source-colony-adjusted means and 95% confidence intervals from the cage-level repeated-measures model. Lines connect model estimates for each dietary treatment across the ascending stimulus series.

### Trehalose levels

The trehalose dataset contained 348 samples, including 10 exact zeros. After accounting for source colony, assay plate, and clustering within cages, we found no clear overall effect of dietary ethanol on trehalose concentration (F_5,3.29_ = 5.11, p = 0.093; Fig. 4). Adjusted estimates varied non-monotonically across treatments (Supplementary Material 1: Table S1.7).

**Figure 4.**
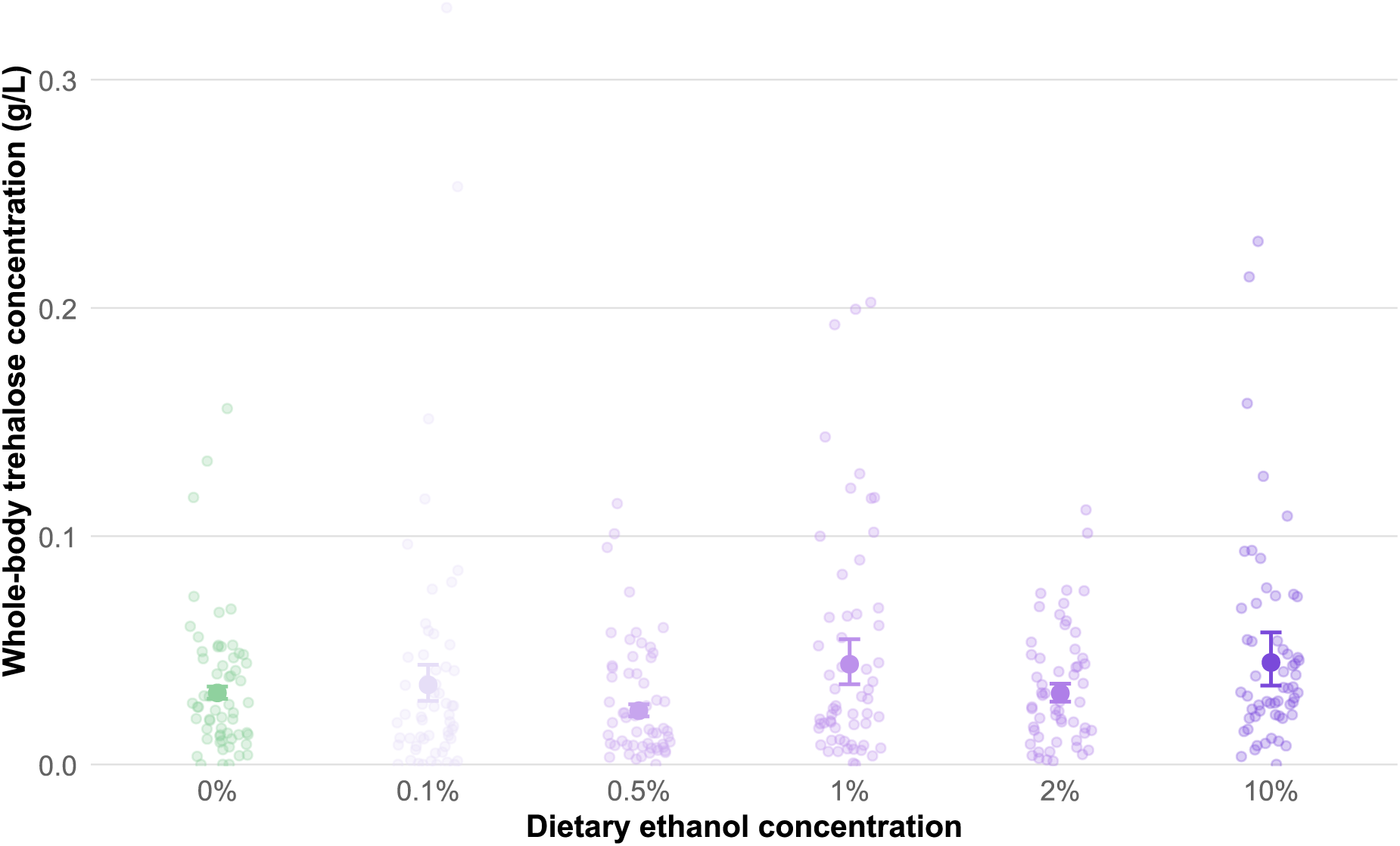
Trehalose concentrations across dietary ethanol treatments. Semi-transparent points represent individual observations. Larger opaque points and error bars show marginal means and 95% confidence intervals from the Tweedie model. Estimates account for source colony and assay plate and use cage-clustered covariance estimates.

### Hemolymph ethanol concentration

Hemolymph ethanol rose sharply with dietary concentration (Fig. 5; Supplementary Material 1: Table S1.8). The probability of an exact zero declined as the dose increased (F_1,9.20_ = 24.1, p < 0.001). Among samples with detectable ethanol, concentration increased with dietary dose (β = 1.398, SE = 0.115, F_1,5.42_ = 147.0, p < 0.001). A one-unit increase in dose corresponded to a 4.05-fold increase in expected positive concentration (95% CI: 3.03–5.41). Overall predicted concentrations rose from 0.0015 g/L in controls to 0.3765 g/L at 10% ethanol.

**Figure 5.**
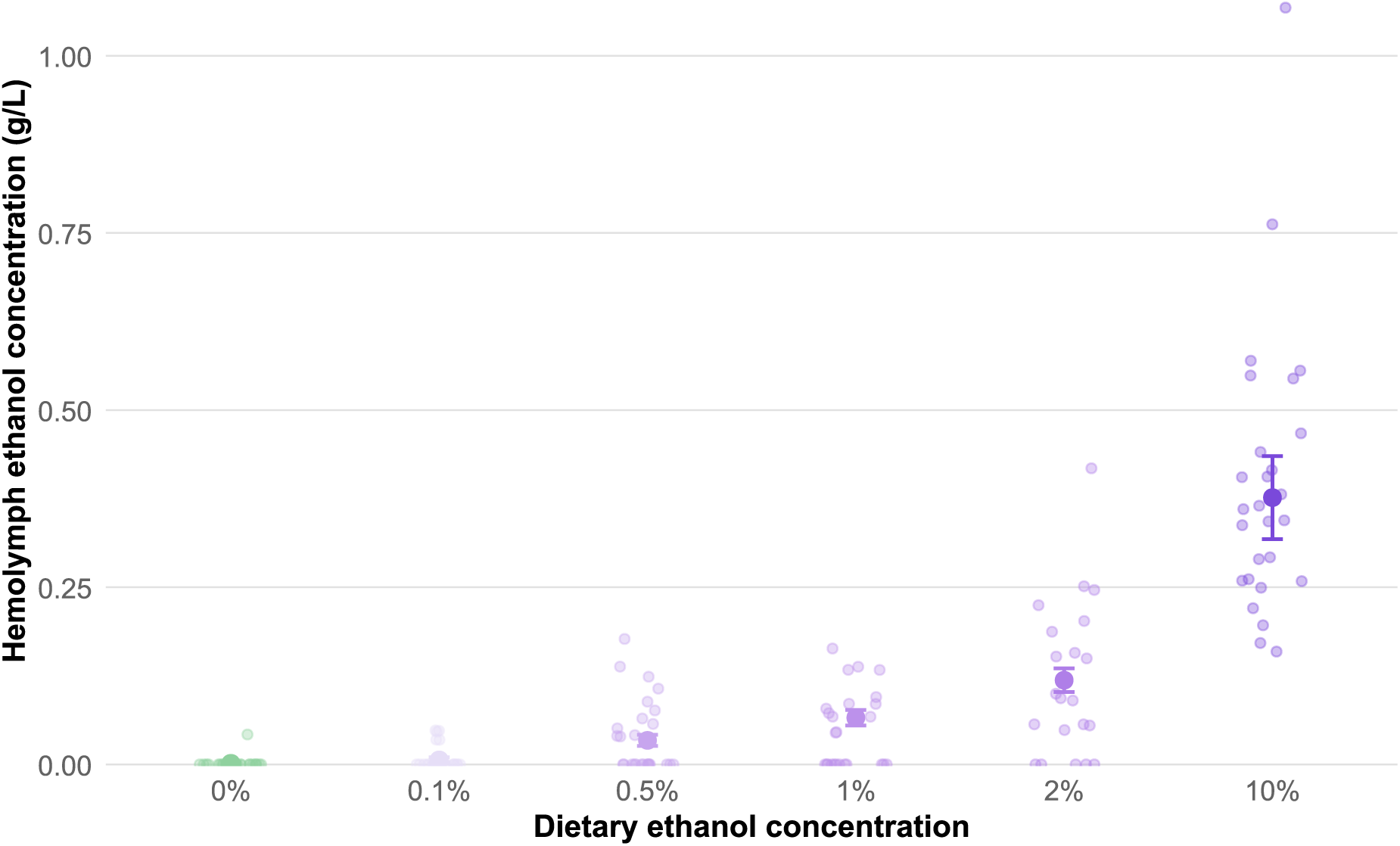
Hemolymph ethanol concentrations across dietary ethanol treatments. Semi-transparent points represent individual observations, including zeros. Larger opaque points and error bars show overall expected concentrations and 95% confidence intervals from the two-part model, averaged across source colonies. These expectations combine the modeled probability of a positive measurement with the expected concentration when ethanol was detected.

## Discussion

Repeated dietary ethanol produced a strong gradient in hemolymph ethanol, but no coordinated hormetic response. Most functional changes were detrimental and appeared at 10%. Bees in this group had the lowest restricted mean survival and estimated intake and showed broad locomotor disruption. Meander was also elevated at 2%. No other ethanol treatment differed clearly from the control. A minor increase in estimated intake at 0.1% was only weakly supported, and dietary treatment did not detectably affect sting extension or trehalose concentration.

The data do not meet the criteria for hormesis under the tested conditions. A hormetic interpretation requires better performance at low exposure than in the untreated control, followed by inhibition at a higher exposure (Agathokleous, 2018; Agathokleous et al., 2018; Agathokleous et al., 2020; Schirrmacher, 2021). Here, no lower-concentration group consistently outperformed the control. The 0.1% group showed a minor trend for greater estimated intake than controls (approximately 9% increase), but bees had no dietary choice, and this result cannot by itself demonstrate preference. Stable performance across several endpoints is still biologically informative, but it is not evidence of benefit. We interpret the pattern as endpoint-specific resilience during eight days of exposure. This interpretation is consistent with reduced attractiveness or stronger toxicity at higher concentrations (Mustard et al., 2019), and the absence of clear impairment after chronic exposure to 0.5-1% ethanol (Ostap-Chec et al., 2025a). More research is needed to determine whether hormetic effects in response to ethanol occur in honeybees in other contexts. In bumblebees, the same broad stress context can produce toxic, neutral, or hormetic effects depending on compound, dose, parameter, and exposure route (Ramanaidu and Cutler, 2013).

The strongest functional effects were locomotor. Nearly all bees began moving, but 10% ethanol reduced the time spent moving, exploration, speed, and distance traveled while increasing meander. At 2%, only meander differed from the control. High dietary ethanol did not simply prevent movement. Bees remained active, but their movement was less sustained, and their paths were shorter and more tortuous. This pattern agrees with acute high-dose studies showing impaired motor coordination (Abramson et al., 2000; Maze et al., 2006). Lower concentrations have produced subtler effects that depend on the assay used (Ahmed et al., 2022; Mixson et al., 2010). Sting extension did not follow the locomotor pattern. Responses increased through the fixed ascending stimulus series, but their overall magnitude and slope did not differ among diets. Thus, chronic dietary ethanol did not measurably alter sting extension, despite evidence that acute ethanol can shift aversive response thresholds (Giannoni-Guzmán et al., 2014). Defensive behavior is strongly context-dependent, with weak or absent ethanol effects in some harnessed-bee assays but increased aggression in free-flying bees under other conditions (Abramson et al., 2000; Abramson et al., 2004b; Ammons and Hunt, 2008). The physiological measurements help separate internal exposure from measurable disruption. Both the likelihood of detecting ethanol and its concentration in positive hemolymph samples rose with dietary dose, as reported in earlier ingestion studies (Bozic et al., 2007; Maze et al., 2006). Whole-body trehalose showed no clear overall treatment effect, but one metabolic marker cannot rule out other physiological costs or compensatory responses. More targeted metabolic measurements may be needed to determine what contributes to ethanol resilience. Comparable ethanol-feeding studies examining trehalose remain scarce (Ostap-Chec et al., 2025a). In that one previous study, bees fed 1% ethanol for two weeks showed a trend towards increased trehalose levels when sampled directly from hemolymph (Ostap-Chec et al., 2025a). Whole-body analyses, as performed here, might have obscured some existing differences between groups.

The ecological interpretation of ethanol exposure is important because ethanol is a widespread fermentation product in sugar-rich resources such as fruits, saps, and nectar, and repeated dietary exposure may have shaped behavioral and metabolic adaptations across taxa (Bowland et al., 2025a; Miler, 2025). Ethanol can also act as a cue associated with fermenting substrates. Ethanol-related attraction or exploratory responses occur, for example, in *Drosophila* and ants (Korczyńska et al., 2023; McKenzie and Parsons, 1972). For honeybees, ethanol is relevant because fermented nectar and overripe fruits may be encountered during foraging. Honeybees detect ethanol-associated cues, voluntarily consume ethanol, and show dose-dependent changes in motor behavior, learning, communication, aggression, trophallaxis, and foraging-related decisions (Abramson et al., 2025). At the same time, ethanol tolerance varies strongly among social Hymenoptera. Oriental hornets tolerate extremely high ethanol concentrations, apparently through rapid metabolism and duplicated alcohol dehydrogenase genes, whereas honeybees are less tolerant under comparable exposure conditions (Bouchebti et al., 2024). Ethanol should therefore be viewed as ecologically relevant for honeybees.

Low-level ethanol appears common in floral nectar. A recent nectar survey found ethanol in at least one sample from 26 of 29 plant species, with concentrations of approximately 0.008-0.056% and species-level values near 0.01-0.02% (Maro et al., 2026a). Higher concentrations can occur, including in bertam palm nectar, where ethanol reached 3.8%, with mean and median values of 0.6% and 0.5%, respectively (Wiens et al., 2008). Fermenting fruits may also expose nectar- and fruit-feeding animals, including bees, to higher ethanol concentrations than those usually measured in floral nectar, although often below 1% (Bowland et al., 2025b; Campbell et al., 2022; Casorso et al., 2023; Dudley, 2002; Dudley, 2004; Eriksson and Nummi, 1982; Maro et al., 2026b). This context is important because honeybees in our experiment remained largely unaffected across the lower and intermediate portion of the tested gradient. Low ethanol exposure may therefore represent a tolerable component of the chemical environment encountered during foraging rather than simply acting as a toxin.

Our findings are relevant for pollinator ecotoxicology because they show that substantial exposure does not necessarily translate directly into measurable impairment. The strength of our design was the combination of a broad ethanol gradient, multiple behavioral and physiological endpoints, and direct hemolymph ethanol measurement. Its main limitation is that it remained cage-based and relatively short-term. Therefore, it cannot determine whether low ethanol concentrations would be harmful, neutral, or beneficial under colony conditions where behavioral choice, social regulation, and resource buffering occur. This limitation is especially relevant because caging itself may alter the honeybee crop microbiota and the background ethanol environment: caged bees can show reduced crop microbial richness, increased community variability, and more frequent detectable crop ethanol, even without access to environmental ethanol sources (Antoł et al., 2025). Future studies should focus more strongly on nectar-realistic ethanol exposure and colony-level consequences. Ethanol should also be tested in combination with pathogens, pesticides, nutritional limitation, or temperature stress, because beneficial effects, if present, may emerge as cross-protection under subsequent challenge rather than as improved performance under ethanol exposure alone. This is particularly relevant given evidence that ethanol access can reduce parasite-driven mortality in some contexts (Ostap-Chec et al., 2025b). Future dose-response experiments should also include several concentrations below 0.1%, because natural nectar ethanol is usually far below the lower bound of many laboratory studies (Abramson et al., 2025; Maro et al., 2026a).

Overall, chronic dietary ethanol produced a marked internal exposure gradient but comparatively limited impairment across the measured endpoints. The clearest costs emerged at 10% ethanol, particularly in food intake and locomotor performance, whereas lower concentrations were generally well tolerated. Evidence for beneficial low-dose stimulation was weak, providing no support for broad ethanol hormesis under these conditions. Instead, the results point to considerable resilience of adult honeybee workers to repeated dietary ethanol exposure. Ethanol in natural resources is an ecologically relevant compound whose consequences depend on concentration, exposure duration, biological context, and the function being measured.

## Acknowledgements

This work was supported by the National Science Centre, Poland [Sonata grant number UMO-2021/43/D/NZ8/01044].

