## Supplementary Material 1 for "Dietary ethanol produces endpoint-specific dose responses in honeybees"

**Table S1.1. Run-adjusted restricted mean survival through day 8 (RMST8) across dietary ethanol treatments.** Estimates were obtained from the cage-level linear model with dietary treatment and experimental run as fixed effects. Standard errors and confidence intervals are based on HC3 heteroscedasticity-consistent covariance estimates. Differences and p-values compare each ethanol treatment with the ethanol-free control using Dunnett adjustment. SE, standard error; LCL, lower confidence limit; UCL, upper confidence limit.

| Dietary ethanol | Estimated RMST8 (days) | Robust SE | LCL | UCL | Difference from control (days) | Dunnett p |
|---|---|---|---|---|---|---|
| 0% | 7.8558 | 0.0702 | 7.7151 | 7.9966 | — | — |
| 0.1% | 7.8258 | 0.0497 | 7.7262 | 7.9255 | −0.0300 | 0.981 |
| 0.5% | 7.7542 | 0.0618 | 7.6302 | 7.8781 | −0.1017 | 0.686 |
| 1% | 7.8933 | 0.0297 | 7.8337 | 7.9529 | 0.0375 | 0.954 |
| 2% | 7.7933 | 0.0362 | 7.7207 | 7.8659 | −0.0625 | 0.847 |
| 10% | 7.5683 | 0.0893 | 7.3894 | 7.7472 | −0.2875 | 0.060 |

**Table S1.2. Estimated mean daily food intake across dietary ethanol treatments.** Estimates were obtained from the cage-level linear model using the complete eight-day exposure period and are adjusted for experimental run. Standard errors and confidence intervals are based on HC3 heteroscedasticity-consistent covariance estimates. Differences and p-values compare each ethanol treatment with the ethanol-free control using Dunnett adjustment. Intake is expressed as mL $bee^{-1}$ $day^{-1}$.

| Dietary ethanol | Mean | Robust SE | 95% CI | Difference from control | Dunnett p |
|---|---|---|---|---|---|
| 0% | 0.03665 | 0.00099 | 0.03466–0.03864 | — | — |
| 0.1% | 0.04004 | 0.00087 | 0.03829–0.04179 | 0.00339 | 0.055 |
| 0.5% | 0.03890 | 0.00109 | 0.03671–0.04109 | 0.00225 | 0.413 |
| 1% | 0.03753 | 0.00063 | 0.03626–0.03879 | 0.00088 | 0.867 |
| 2% | 0.03843 | 0.00047 | 0.03749–0.03938 | 0.00178 | 0.357 |
| 10% | 0.02842 | 0.00109 | 0.02624–0.03060 | −0.00823 | <0.001 |

**Table S1.3. Locomotor responses across dietary ethanol treatments.** Overall treatment tests are from the cage-level models including source colony as a fixed factor; p-values were adjusted across the five locomotor outcomes using the Holm method. Treatment estimates are source-colony-adjusted marginal means with 95% confidence intervals, back-transformed to the original measurement scale. Treatment-specific p-values compare each ethanol treatment with the ethanol-free control using Dunnett adjustment.

**A. Overall treatment tests**

| Endpoint | Cage-level F | Primary p | Holm p |
|---|---|---|---|
| Proportion of time moving | 7.84 | 0.0031 | 0.0097 |
| Relative exploration | 8.36 | 0.0024 | 0.0097 |
| Average speed while moving | 5.04 | 0.0145 | 0.0145 |

| Endpoint | Cage-level F | Primary p | Holm p |
|---|---|---|---|
| Distance traveled while moving | 7.46 | 0.0037 | 0.0097 |
| Meander while moving | 11.04 | <0.001 | 0.0041 |

### B. Treatment estimates and comparisons with control

| Endpoint | Dietary ethanol | Adjusted estimate [95% CI] | Dunnett p |
|---|---|---|---|
| Proportion of time moving | 0% | 0.788 [0.725, 0.839] | — |
| Proportion of time moving | 0.1% | 0.792 [0.730, 0.842] | 0.999 |
| Proportion of time moving | 0.5% | 0.832 [0.779, 0.874] | 0.553 |
| Proportion of time moving | 1% | 0.766 [0.700, 0.822] | 0.933 |
| Proportion of time moving | 2% | 0.712 [0.638, 0.777] | 0.290 |
| Proportion of time moving | 10% | 0.594 [0.510, 0.672] | 0.006 |
| Relative exploration | 0% | 0.741 [0.664, 0.805] | — |
| Relative exploration | 0.1% | 0.677 [0.592, 0.752] | 0.558 |
| Relative exploration | 0.5% | 0.754 [0.680, 0.816] | 0.987 |
| Relative exploration | 1% | 0.670 [0.584, 0.745] | 0.480 |
| Relative exploration | 2% | 0.613 [0.523, 0.696] | 0.111 |
| Relative exploration | 10% | 0.452 [0.363, 0.544] | 0.001 |
| Average speed while moving | 0% | 2.686 [2.485, 2.904] | — |
| Average speed while moving | 0.1% | 2.642 [2.443, 2.856] | 0.982 |
| Average speed while moving | 0.5% | 2.894 [2.676, 3.128] | 0.465 |
| Average speed while moving | 1% | 2.594 [2.400, 2.805] | 0.886 |
| Average speed while moving | 2% | 2.532 [2.342, 2.737] | 0.636 |
| Average speed while moving | 10% | 2.279 [2.108, 2.464] | 0.030 |
| Distance traveled while moving | 0% | 1062 [855, 1319] | — |
| Distance traveled while moving | 0.1% | 1071 [862, 1329] | 1.000 |
| Distance traveled while moving | 0.5% | 1313 [1058, 1631] | 0.441 |
| Distance traveled while moving | 1% | 976 [786, 1211] | 0.917 |
| Distance traveled while moving | 2% | 871 [701, 1081] | 0.493 |

| Endpoint | Dietary ethanol | Adjusted estimate [95% CI] | Dunnett p |
|---|---|---|---|
| Distance traveled while moving | 10% | 599 [483, 744] | 0.008 |
| Meander while moving | 0% | 333.2 [311.1, 356.9] | — |
| Meander while moving | 0.1% | 362.4 [338.3, 388.1] | 0.271 |
| Meander while moving | 0.5% | 318.5 [297.4, 341.1] | 0.724 |
| Meander while moving | 1% | 353.8 [330.4, 379.0] | 0.533 |
| Meander while moving | 2% | 390.9 [365.0, 418.7] | 0.018 |
| Meander while moving | 10% | 420.8 [392.9, 450.7] | 0.001 |

**Table S1.4. Model-estimated sting extension and voltage-response slopes across dietary ethanol treatments.** Estimates are derived from the cage-level repeated-measures model with dietary treatment, voltage, their interaction, and source colony as fixed effects, and treatment cage fitted with uncorrelated random intercept and voltage slope. Sting-extension estimates refer to the midpoint of the stimulus series (4 V). Voltage slopes represent the estimated change in sting extension per 1 V increase. Differences from the ethanol-free control and associated p-values are Dunnett-adjusted. Values in brackets are 95% confidence intervals.

| Dietary ethanol | Sting extension at 4 V, mm [95% CI] | Difference from control, mm [95% CI] | Dunnett p | Voltage slope, mm $V^{-1}$ [95% CI] | Slope difference from control [95% CI] | Dunnett p |
|---|---|---|---|---|---|---|
| 0% | 0.741 [0.602, 0.880] | — | — | 0.097 [0.049, 0.146] | — | — |
| 0.1% | 0.656 [0.517, 0.795] | −0.085 [−0.352, 0.181] | 0.763 | 0.080 [0.032, 0.129] | −0.017 [−0.109, 0.075] | 0.938 |
| 0.5% | 0.825 [0.686, 0.964] | 0.083 [−0.183, 0.350] | 0.775 | 0.119 [0.071, 0.167] | 0.022 [−0.070, 0.114] | 0.891 |
| 1% | 0.684 [0.545, 0.824] | −0.057 [−0.323, 0.210] | 0.908 | 0.123 [0.074, 0.171] | 0.025 [−0.067, 0.117] | 0.841 |
| 2% | 0.756 [0.617, 0.895] | 0.014 [−0.252, 0.281] | 0.998 | 0.114 [0.066, 0.163] | 0.017 [−0.075, 0.109] | 0.942 |
| 10% | 0.864 [0.724, 1.003] | 0.122 [−0.144, 0.389] | 0.528 | 0.117 [0.068, 0.165] | 0.019 [−0.073, 0.111] | 0.918 |

**Table S1.5. Model-estimated sting extension across the seven electric stimuli.** Values are source-colony-adjusted estimated mean sting extension in mm, with 95% confidence intervals in brackets, from the cage-level repeated-measures model used for Fig. 3. Estimates include zero responses.

| Dietary ethanol | 1 V | 2 V | 3 V | 4 V | 5 V | 6 V | 7 V |
|---|---|---|---|---|---|---|---|
| 0% | 0.449 [0.260, 0.639] | 0.547 [0.386, 0.707] | 0.644 [0.500, 0.788] | 0.741 [0.602, 0.880] | 0.839 [0.695, 0.982] | 0.936 [0.775, 1.097] | 1.033 [0.844, 1.223] |
| 0.1% | 0.416 [0.226, 0.605] | 0.496 [0.335, 0.656] | 0.576 [0.432, 0.720] | 0.656 [0.517, 0.795] | 0.736 [0.592, 0.880] | 0.816 [0.656, 0.977] | 0.896 [0.707, 1.086] |
| 0.5% | 0.468 [0.278, 0.657] | 0.587 [0.426, 0.747] | 0.706 [0.562, 0.849] | 0.825 [0.686, 0.964] | 0.944 [0.800, 1.088] | 1.063 [0.902, 1.223] | 1.182 [0.992, 1.371] |
| 1% | 0.316 [0.127, 0.506] | 0.439 [0.278, 0.599] | 0.562 [0.418, 0.705] | 0.684 [0.545, 0.824] | 0.807 [0.663, 0.951] | 0.930 [0.769, 1.091] | 1.053 [0.863, 1.242] |
| 2% | 0.413 [0.224, 0.603] | 0.527 [0.367, 0.688] | 0.641 [0.498, 0.785] | 0.756 [0.617, 0.895] | 0.870 [0.726, 1.014] | 0.984 [0.823, 1.145] | 1.098 [0.909, 1.288] |
| 10% | 0.514 [0.324, 0.703] | 0.630 [0.470, 0.791] | 0.747 [0.603, 0.891] | 0.864 [0.724, 1.003] | 0.980 [0.837, 1.124] | 1.097 [0.936, 1.258] | 1.214 [1.024, 1.403] |

**Table S1.6. Pre-stimulus sting extension across dietary ethanol treatments.** Pre-stimulus measurements were analyzed as supplementary control. Because dietary treatment was assigned at the cage level, sting extension measured immediately before each stimulus was averaged across all bees and all seven positions in the stimulus sequence within each treatment cage, including zero values when no extension was visible. This yielded one response for each of the 18 treatment cages. Cage-level mean extension was analyzed using a linear model with dietary treatment and source colony as fixed effects. Statistical inference used HC3 heteroscedasticity-consistent covariance estimates; the overall treatment effect was assessed with an HC3-robust joint F-test, and each ethanol treatment was compared with the ethanol-free control using Dunnett adjustment. Mean pre-stimulus extension did not differ among dietary treatments ($F_{5,10} = 1.05$, $p = 0.439$), and none of the ethanol treatments differed from the control after Dunnett adjustment (all $p \geq 0.411$). Adjusted means and 95% confidence intervals are model-based estimates controlling for source colony. Mean cage proportion positive is the descriptive mean across the three treatment cages of the proportion of pre-stimulus measurements showing visible sting extension. Sequence position was not included in the reported treatment analysis because each pre-stimulus measurement preceded the corresponding electrical stimulus, and position within the assay was inseparable from elapsed time and cumulative exposure to preceding shocks.

| Dietary ethanol | Adjusted mean extension, mm [95% CI] | Mean cage proportion positive | Difference from control, mm [95% CI] | Dunnett p |
|---|---|---|---|---|
| 0% | 0.074 [−0.044, 0.193] | 0.124 | — | — |
| 0.1% | 0.192 [0.075, 0.310] | 0.295 | 0.118 [−0.108, 0.344] | 0.426 |
| 0.5% | 0.256 [−0.070, 0.583] | 0.405 | 0.182 [−0.289, 0.653] | 0.653 |
| 1% | 0.132 [0.120, 0.144] | 0.229 | 0.057 [−0.104, 0.219] | 0.707 |
| 2% | 0.198 [0.012, 0.385] | 0.343 | 0.124 [−0.175, 0.423] | 0.605 |
| 10% | 0.226 [0.052, 0.400] | 0.381 | 0.152 [−0.133, 0.436] | 0.411 |

**Table S1.7. Model-estimated marginal trehalose concentrations across dietary ethanol treatments.** Estimates are presented on the response scale in g/L and are adjusted for source colony and assay plate. Standard errors and 95% confidence intervals were calculated using the cage-clustered CR2 covariance matrix. EMMean, estimated marginal mean; SE, standard error; LCL, lower confidence limit; UCL, upper confidence limit.

| Dietary ethanol | EMMean (g/L) | SE | LCL | UCL |
|---|---|---|---|---|
| 0% | 0.0313 | 0.0037 | 0.0248 | 0.0395 |
| 0.1% | 0.0348 | 0.0041 | 0.0277 | 0.0438 |
| 0.5% | 0.0235 | 0.0029 | 0.0184 | 0.0300 |
| 1% | 0.0439 | 0.0049 | 0.0353 | 0.0546 |
| 2% | 0.0312 | 0.0038 | 0.0245 | 0.0396 |
| 10% | 0.0447 | 0.0049 | 0.0360 | 0.0554 |

**Table S1.8. Observed zero measurements and predicted overall hemolymph ethanol concentrations across dietary treatments.** Predictions were obtained from the two-part model and averaged across source colonies. Overall predictions combine the predicted probability of positive measurement with the predicted ethanol concentration conditional on detection. Confidence intervals are model-based 95% confidence intervals. Overall predicted concentrations are provided for presentation; statistical inference was based on the separate cage-clustered CR2 tests for zero probability and positive ethanol concentration.

| Dietary ethanol | Observations | Exact zeros, n (%) | Predicted concentration (g/L) | 95% CI |
|---|---|---|---|---|
| 0% | 23 | 22 (95.7%) | 0.0015 | 0.00001–0.0030 |

| Dietary ethanol | Observations | Exact zeros, n (%) | Predicted concentration (g/L) | 95% CI |
|---|---|---|---|---|
| 0.1% | 26 | 21 (80.8%) | 0.0063 | 0.0025–0.0101 |
| 0.5% | 27 | 15 (55.6%) | 0.0339 | 0.0260–0.0417 |
| 1% | 25 | 12 (48.0%) | 0.0658 | 0.0549–0.0767 |
| 2% | 21 | 5 (23.8%) | 0.1188 | 0.1021–0.1355 |
| 10% | 27 | 0 (0.0%) | 0.3765 | 0.3180–0.4350 |
